# Temporal inhibition of autophagy reveals segmental reversal of aging with increased cancer risk

**DOI:** 10.1101/528984

**Authors:** Liam D Cassidy, Andrew RJ Young, Christopher NJ Young, Elizabeth J Soilleux, Edward Fielder, Bettina M Weigand, Rebecca Brais, Kimberley A Wiggins, Murray CH Clarke, Diana Jurk, Joao F Passos, Masashi Narita

## Abstract

Autophagy is an important cellular degradation pathway with a central role in metabolism as well as basic quality control, two processes inextricably linked to aging. A decrease in autophagy is associated with increasing age, yet it is unknown if this is causal in the aging process, and whether autophagy restoration can counteract these aging effects. Here we demonstrate that systemic autophagy inhibition induces the premature acquisition of age-associated phenotypes and pathologies in mammals. Remarkably, autophagy restoration provides a near complete recovery of morbidity and a significant extension of lifespan, however, at the molecular level this rescue appears incomplete. Importantly autophagy-restored mice still succumb earlier due to an increase in spontaneous tumor formation. Thus our data suggest that chronic autophagy inhibition confers an irreversible increase in cancer risk and uncovers a biphasic role of autophagy in cancer development being both tumor suppressive and oncogenic, sequentially.

## Main Text

Physiological aging is a complex and multifaceted process associated with the development of a wide array of degenerative disease states. While there is no accepted singular underlying mechanism of aging, a combination of genetic, environmental and metabolic factors have been shown to alter the aging process^1–3^. As such, lifestyle and pharmacological regimens have been proposed that may offer health- and or life-span benefits^4–6^. However, despite chronological aging representing the greatest risk factor for pathological conditions as diverse as neurodegeneration, cancer, and cardiovascular disease, there is a paucity of genetic mammalian models that allow for dynamic modulation of key processes in mammalian aging.

Autophagy is an evolutionarily conserved bulk cellular degradation system that functions to breakdown and recycle a wide array of cytoplasmic components from lipids, proteins and inclusion bodies, to whole organelles (e.g. mitochondria). Importantly a reduction in autophagic flux (the rate at which autophagosomes form and breakdown cellular contents) is associated with increasing age in mammals^7^. Evidence from lower organisms suggests that autophagy inhibition can negate the positive-effects of regimens that extend lifespan, such as calorie restriction, rapamycin supplementation, and mutations in insulin signalling pathways^8–10^.

In mice, the constitutive promotion of autophagy throughout lifetime has been shown to extend health- and life-span in mammalian models^11, 12^. These studies have provided hitherto missing evidence that autophagic flux can impact on mammalian longevity and supports the notion that the pharmacological promotion of autophagy may extend health-, and potentially life-span, in humans. However, whether a reduction in autophagy is sufficient to induce phenotypes associated with aging, and whether these effects can be reversed by restoring autophagy has to date not been addressed.

Considering that the therapeutic window for pharmacological intervention to counteract aging, and age-related diseases, will be later in life (as opposed to from conception), after autophagic flux has declined, it is critical to understand how the temporal modulation (inhibition and restoration) of autophagy may impact on longevity and health.

To address these questions, we use two doxycycline (dox) inducible shRNA mouse models that target the essential autophagy gene Atg5 (Atg5i mice) to demonstrate that autophagy inhibition in young adult mice is able to drive the development of aging-like phenotypes and reduce longevity. Importantly we confirm that the restoration of autophagy is associated with a substantial restoration of health- and life-span, however this recovery is incomplete. Notably the degree of recovery is segmental, being dependent on both the tissue and metric analysed. A striking consequence of this incomplete restoration is that autophagy restored mice succumb to spontaneous tumor formation earlier and at an increased frequency than control mice, a phenotype not observed during autophagy inhibition alone. As such our studies indicate that despite the significant benefit, autophagy reactivation may also promote tumorigenesis in advanced ageing context.

## Reduced lifespan in Atg5i mice

Previously, we have reported the development of a highly efficient dox inducible shRNA mouse model targeting Atg5 (Atg5i)^13^ that phenocopies tissue-specific Atg5 knockout (KO) mice. These mice lack brain expression of the shRNA and as such do not suffer from the lethal neurotoxic effects that characterise systemic autophagy knockout mice^14, 15^, and enable us to perform longitudinal studies that were previously unachievable *in vivo.*

A common caveat of many mouse models is that genetic manipulations are often present during embryogenesis. Thus, any phenotypes that manifest are a combination of both developmental and tissue homeostasis effects. To avoid the generation of these compound effects, Atg5i mice were aged until eight-weeks (young adults) before being transferred to a dox-containing diet and followed to assess overall survival. Atg5i mice on long-term dox (LT-Atg5i) had a median survival of ~six months on dox (Male 185 days; Female 207 days on dox) with no apparent sex bias (Fig. 1a-c and Supplementary Fig. 1a).

**Figure 1:**
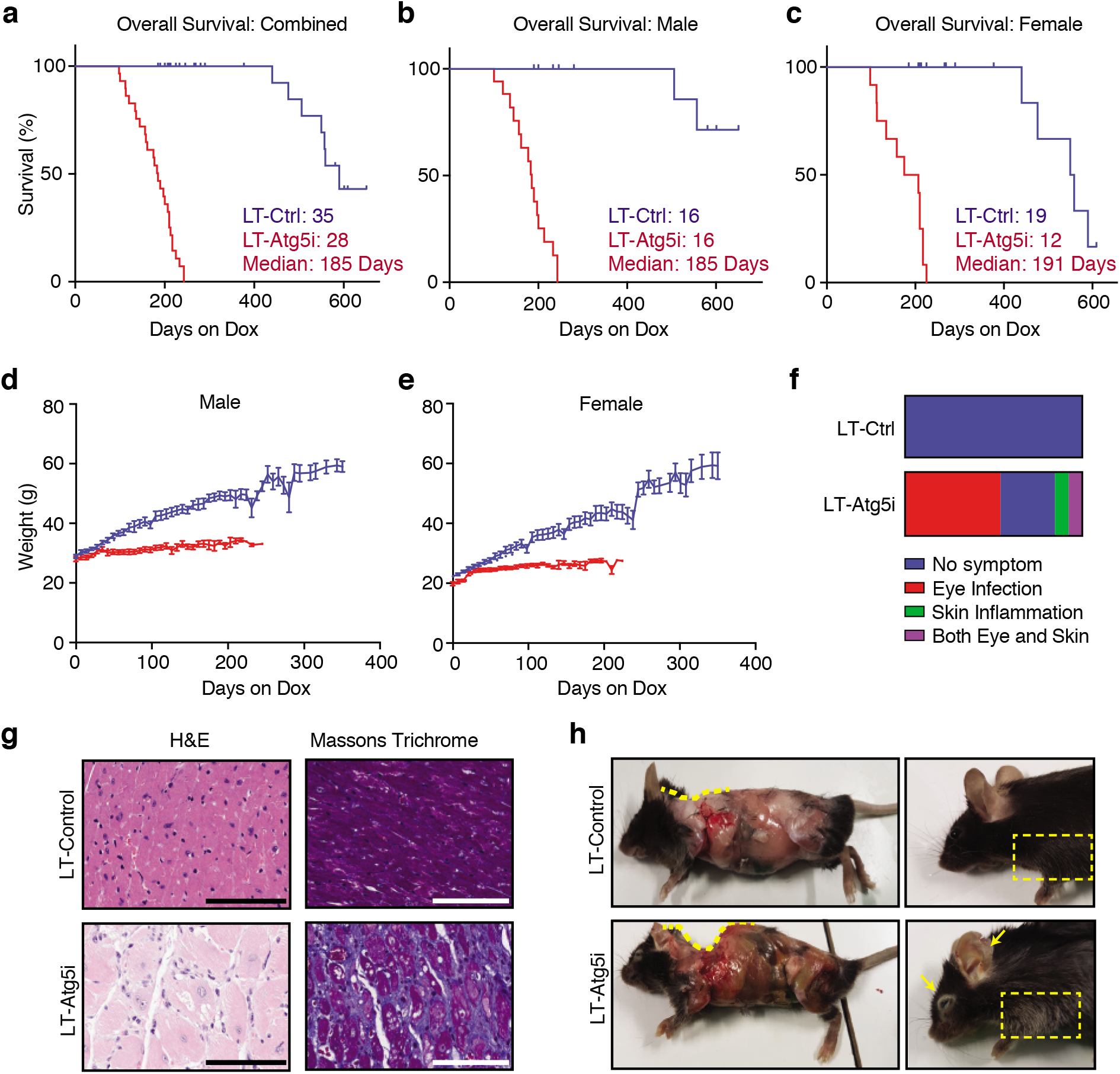
Autophagy inhibition decreases lifespan. **a-c**, LT-Atg5i mice on dox continuously from two months old display a reduced lifespan in comparison to LT-Control as shown in survival graphs for **(a)** combined (p<0.0001), **(b)** male (p<0.0001), **(c)** female (p<0.0001) (Mantel-Cox test). Median survival (days on dox) and mice per group are indicated. **d-e**, During this period LT-Atg5i mice also display a reduced weight gain in both **(d)** male and **(e)** female cohorts. **f**, LT-Atg5i mice also display an increased frequency of skin inflammation and eye infections in comparison to age-matched LT-Control mice. **g**, Cardiac fibrosis was also evident in LT-Atg5i mice. Representative images of H&E and Massons Trichrome are shown. Scale bars,100 μm. h, Age-matched skinned mice. LT-Atg5i mice show kyphosis (yellow dotted line traces the arch of the spine). They often displayed premature greying (dotted rectangle). Arrows indicate the presence of inflammation.

In comparison to littermate controls, LT-Atg5i mice experienced a progressive deterioration, initially presenting with a reduction in coat condition within the first few weeks and a reduction in weight gain that became more pronounced over the life of the animal (Fig. 1d, e and Supplementary Fig. 1b). The majority of mice eventually succumbed to a general morbidity characterised by lethargy, piloerection, and a decrease in body condition, wherein they have to be sacrificed. As previously described with naturally aged colonies^16^, LT-Atg5i mice also appeared susceptible to eye infections and ulcerative dermatitis, the later being primarily localised to the ears and neck and ranging from mild to severe (Fig. 1f and Supplementary Fig. 1c, respectively).

A singular cause of death in LT-Atg5i mice is difficult to determine and it is most likely of multifactorial aetiology across the cohort. At necropsy, all mice displayed hepatomegaly and splenomegaly in comparison to age and sex matched controls, consistent with phenotypes associated with tissue specific knockout mice^17–19^. Elevated serum ALT and reduced levels of serum albumin were present throughout dox administration of Atg5i mice, yet were altered further at the time of death only in a subset of samples (Supplementary Fig. 1f, g, yellow circles). Consistent with this, an increase in serum bilirubin levels was only observed at the time of death within this same subset of mice (Supplementary Fig. 1h, yellow circle). These data suggest that severe liver failure occurs in only a fraction.

Interestingly serum creatinine levels, a marker of kidney function, also displayed an increase only in a different subset of LT-Atg5i mice at the time of death, although they were not generally high during dox administration (Supplementary Fig. 2a). Loss of autophagy also correlated with a general thickening of the basement membrane and the presence of sclerotic (Supplementary Fig. 2b) and enlarged glomeruli (Supplementary Fig. 2c, d) in comparison to age-matched tissue samples, indicative of degenerative kidney disorder. LT-Atg5i mice also stained positively for the build-up of toxic amyloid proteins and the autophagy adaptor protein p62/Sqstm1, a condition normally associated with advancing age in humans (Supplementary Fig. 2e, f). These data suggest that, similar to the liver, systemic autophagy defect causes age-related degenerative alterations in kidney, yet only a distinct subset progresses to renal failure on death. In addition to this stochastic development of organ failure, LT-Atg5i mice universally presented with cardiomyopathy (Supplementary Fig. 2g). Histological examination highlighted the presence of enlarged, degenerate and vacuolated cardiomyocytes, in addition to the presence of cardiac fibrosis (Fig. 1g).

Together, our data suggest that, despite the stereotypic premature death, LT-Atg5i mice suffered from a heterogeneous set of tissue degenerative disorders that appear to have contributed to an increase in mortality. Of note, there was no evidence of overt tumor development in these mice.

## Autophagy inhibition is associated with accelerated aging

All LT-Atg5i mice displayed evidence of kyphosis after four months of dox treatment that became progressively more pronounced as the animals aged until death, whilst 16/28 LT-Atg5i mice displayed evidence of premature greying to varying degrees (Fig. 1h). Furthermore LT-Atg5i mice displayed evidence of extramedullary hematopoiesis (Fig. 2a) and immune aggregations, commonly seen in aged mouse colonies, were also found in the liver, lungs and kidneys but were generally absent in age matched controls, although incidence of these increased in frequency with increasing age (Supplementary Fig. 3a).

**Figure 2:**
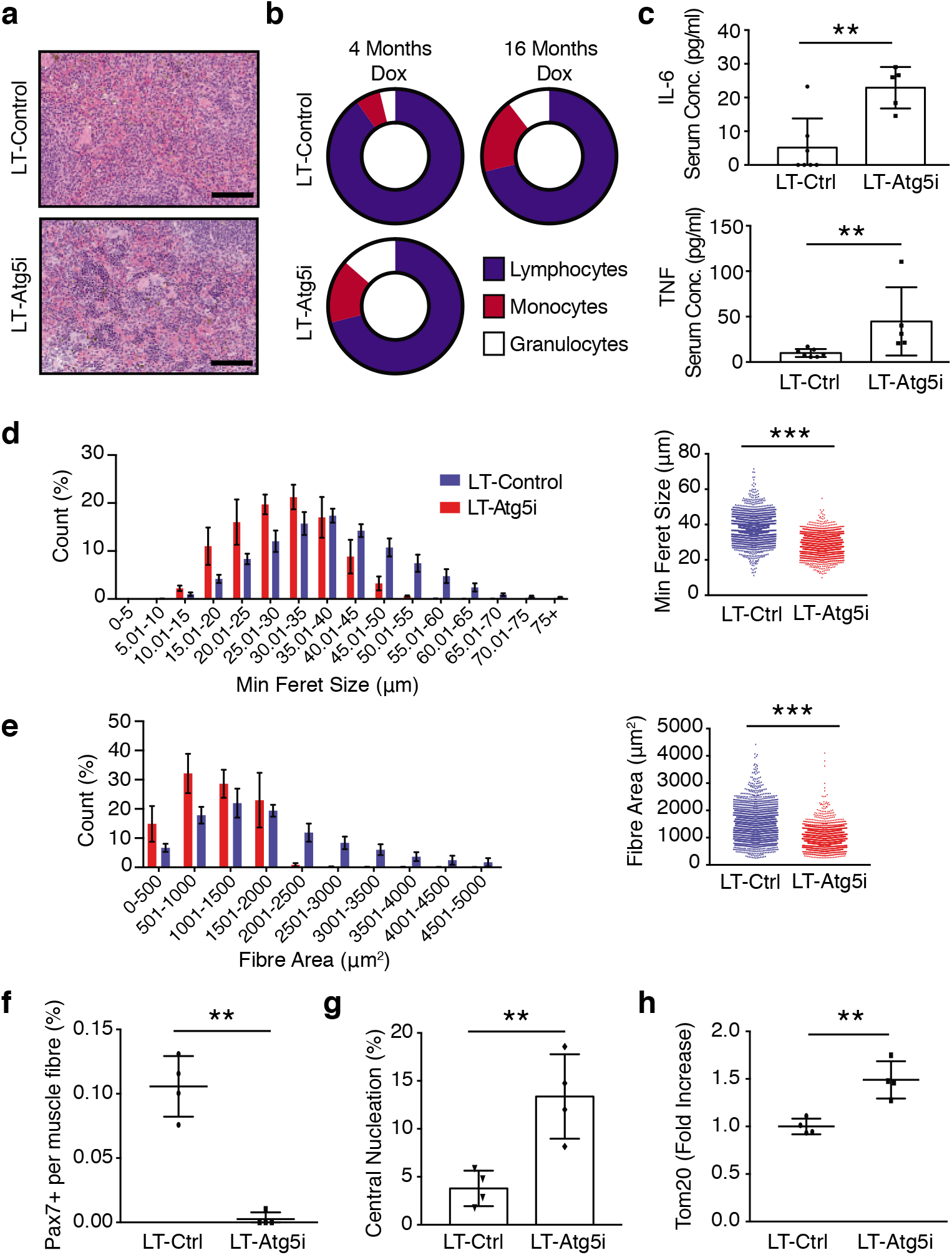
LT-Atg5i mice present with accelerated aging phenotypes. **a**, Extramedullary haematopoiesis is present in the spleens of LT-Atg5i mice in comparison to age-matched controls. Scale bars,100 μm. **b**, Composition of the peripheral immune system in LT-Atg5i mice is reminiscent of old control mice. (n=5-6 mice per group). **c**, Six-month-old LT-Atg5i mice (four months dox treatment) displayed increased serum levels of IL-6 and TNF (LT-Atg5i n=5, LT-Ctrl n=8; Mann Whitney Test). **d-h**, LT-Atg5i mice display alterations in skeletal muscle after six-months of dox treatment. **(d)** LT-Atg5i mice display a significant difference in minimum feret size (n= 4 R-Ctrl and 4 R-Atg5i, Mann Whitney test) and **(e)** cross-sectional area (n= 4 R-Ctrl and 4 R-Atg5i, Mann Whitney test). LT-Atg5i mice also display a decrease in Pax7 nuclear positivity per fibre **(f)**, an increase in central nucleation **(g)**, and positivity for the mitochondrial marker TOM20 **(h)**, as determined by tissue immunofluorescence (unpaired two-tailed Welches t-test; n= 4 R-Ctrl and 4 R-Atg5i). Error bars indicate standard deviations. *p<0.05; **p<0.01, ***p<0.001

As previously described in hematopoietic Atg5 KO mice, LT-Atg5i mice also displayed an increase in cellularity of the peripheral immune system^18, 20^ (Supplementary Fig. 3b) with a myeloid skewing (Fig. 2b) reminiscent of age-associated chronic inflammation. This ‘inflamm-aging’ phenotype was further supported by an increase in serum TNF and IL-6 in LT-Atg5i mice in comparison to control (Fig. 2c).

Skeletal muscle exhibits an age-related decline and autophagy has been reported to be required for the maintenance of Pax7 positive satellite cells (myogenic precursors)^21^. In accordance, LT-Atg5i mice displayed evidence of skeletal muscle degeneration with the presence of smaller fibres, a reduction in the population of Pax7 positive satellite cells, and an increase in central nucleation in comparison to age-matched littermate control mice (Fig. 2d-g). Central nucleation represents muscle fibre regeneration after acute muscle injury but an increase in basal frequency of centrally nucleated myofibres is also a sign of sarcopenia at geriatric age both in mice and human^22^. Additionally, LT-Atg5i muscle fibres displayed increased staining positivity for the mitochondrial marker TOM20 indicative of increased mitochondrial mass and a reduction in autophagy mediated turnover (Fig. 2h).

The accumulation of senescent cells is considered a key marker of chronological aging. Autophagy has been reported to have context dependent and sometimes opposing roles during cellular senescence: typically basal autophagy is considered to promote fitness and its loss may promote senescence, whereas in oncogene-induced senescence, autophagy may be important for the establishment of senescent phenotypes^23–26^. To determine if the systemic loss of basal autophagy is sufficient to drive the establishment of cellular senescence *in vivo* we performed western blotting across a number of tissues from 4-month dox treated LT-Atg5i mice and found an increased staining pattern for key senescence markers (i.e. p16, p21, and p53) (Fig. 3a-c and Supplementary Fig. 3c). Additionally, whole mount senescence-associated beta-galactosidase staining from 6-month treated livers highlighted a marked increase in staining patterns in comparison to LT-Control mice (Fig. 3d). Histologically, nuclear accumulation of p21 was also evident, particularly in hepatocytes with enlarged morphology (Fig. 3d). Furthermore LT-Atg5i mice display a significant increase in both the abundance and frequency of telomere-associated γ-H2AX foci (TAF), which have been shown to correlate with senescence, increasing age and mitochondrial dysfunction (Fig. 3e, f)^27–29^. These data reinforce the age acceleration upon systemic autophagy reduction.

**Figure 3:**
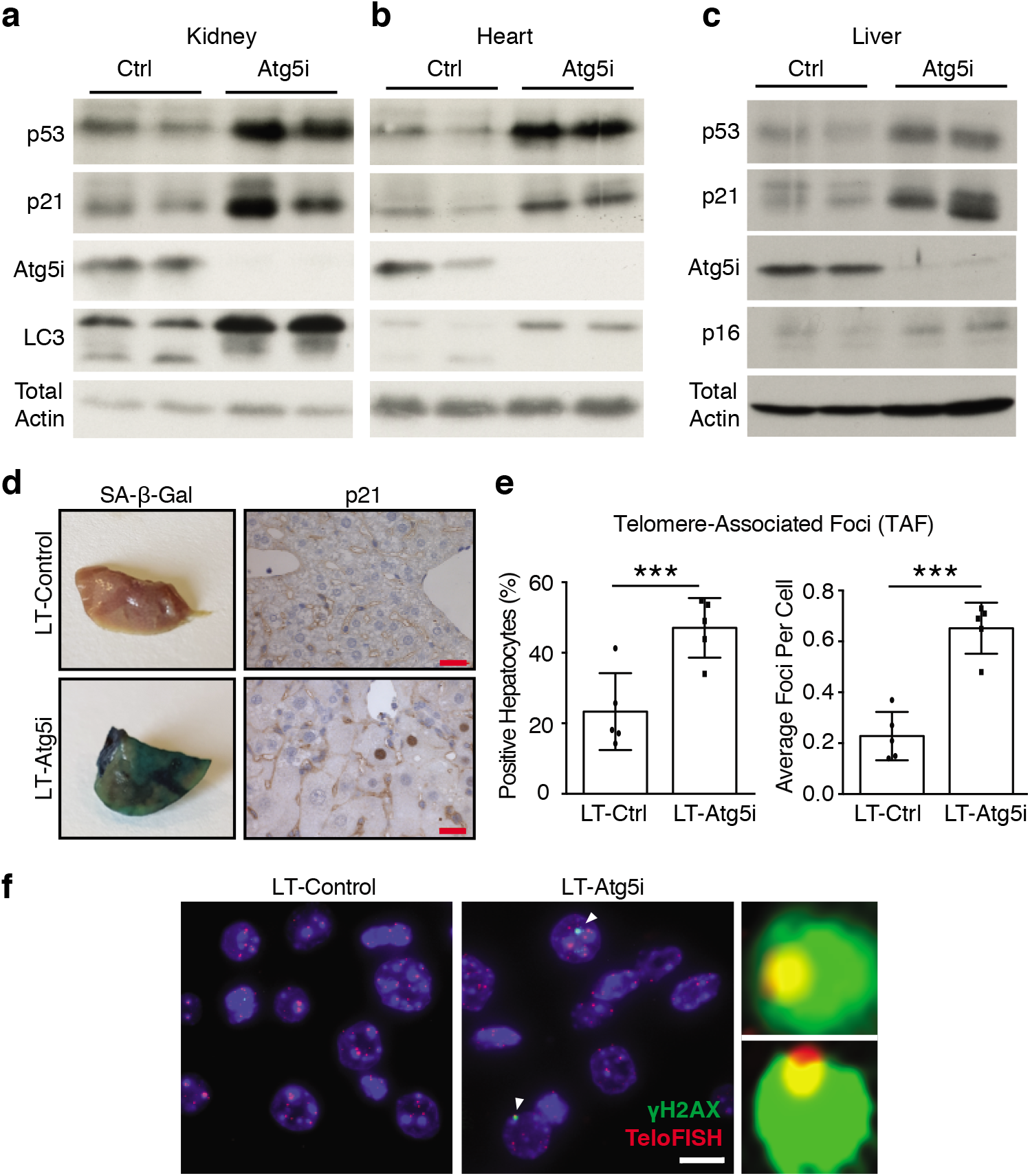
Autophagy inhibition drives senescence in vivo. **a-d**, Markers of senescence can also be seen across multiple tissues in our LT-Atg5i cohorts treated with dox for four months including in **(a)** kidney, **(b)** heart, and **(c)** liver. **(d)** LT-Atg5i livers stain positively for senescence associated β-galactosidase and p21 unlike age-matched control mice (scale bar, 25 μm). **e**, Six-month doxycycline treated LT-Atg5i livers display an increase in the frequency and abundance of γ-H2AX at telomeres, a marker associated with increasing chronological age (unpaired two-tailed t-test; n=5). **f**, A representative example image shown. Arrowheads point to TAF that are magnified on the right of the image. Scale bar, 10 μm. Error bars indicate standard deviation ***p<0.001

Of note, similar gross phenotypic results were also seen in mice with a second hairpin targeting Atg5 (LT-Atg5i_2). LT-Atg5i_2 mice display evidence of premature aging-like phenotypes (Supplementary Fig. 4a-c), however the appearance of these phenotypes is delayed in comparison to LT-Atg5i mice, seemingly due to a hypomorphic reduction in Atg5. Accordingly, these mice displayed the accumulation of p62/Sqstm1 and LC3 in multiple tissues but at lower levels in comparison to LT-Atg5i mice, and did not display phenotypes associated with complete Atg5 knockout mice, includin hepatomegaly and splenomegaly (Supplementary Fig. 4d-f). These findings in particular are important as they establish that the reduction in longevity and presence of aging phenotypes is not dependent on the hepatomegaly and splenomegaly phenotypes encountered in the original LT-Atg5i mouse strain with the highest degree of autophagy inhibition.

Combined these data support a role for basal autophagy in maintaining tissue and organismal homeostasis and provide evidence that causally links autophagy inhibition to the induction of aging-like phenotypes in mammals.

## Autophagy Restoration Partially Reverses Accelerated Aging-like Phenotypes

We next sought to determine whether autophagy restoration alone is able to reverse the aging-like phenotypes by removing dox from the diet. Eight-week old Atg5i and control mice treated with dox for four months, the point at which they universally presented with kyphosis, were switched back to a diet absent of dox leading to a restoration in Atg5 levels and autophagy (termed R-Atg5i cohort)^13^. Interestingly the senescence marker p21 remained elevated across a number of tissues 2 months post dox removal (Fig. 4a, b).

**Figure 4:**
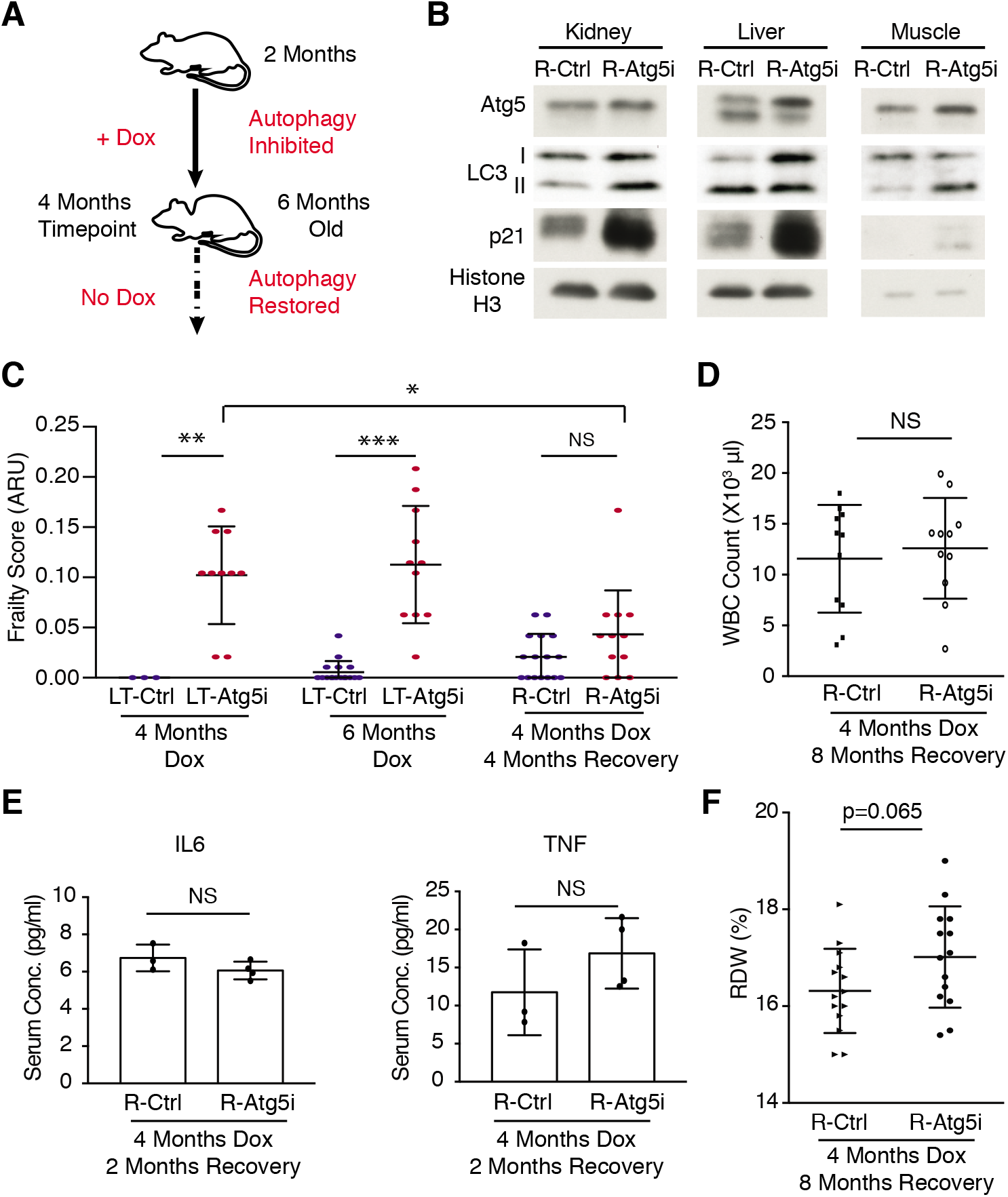
Restoration of autophagy partially restores health-span. **a**, Schematic of R-Atg5i study. Briefly two-month old mice are given dox to induce Atg5 downregulation for four months at which point they exhibit ageing-like phenotypes. Dox is then removed and autophagy restored. **b**, Tissues from R-Atg5i mice with autophagy restored for two months display evidence of ATG5 protein and autophagy restoration, yet still stain positively for markers of senescence. **c**, Atg5i mice on dox for four months and six months display increase frailty scores in comparison to controls (ARU, arbitrary units). While R-Atg5i mice where autophagy has been restored for four months, display a recovery (Two-way ANOVA with Tukey’s correction for all comparisons, n=3-16). **d**, Whole blood cell counts from R-Atg5i mice display no difference in comparison to age matched R-Control mice (unpaired two-tailed t-test; n=11 per group). **e**, Inflammatory serum cytokines IL6 and TNF are equivalent in R-Atg5i and R-Control mice two-months post dox removal (Mann Whitney test; n= 3 R-Ctrl and 4 R-Atg5i). **f**, Red blood cell distribution width (RDW) is altered in aged autophagy-restored cohorts (four months dox, eight months restoration) (unpaired two-tailed t-test; n=14 per group). Error bars indicate standard deviation; NS denotes not significant. *p<0.05; **p<0.01, ***p<0.001.

An increase in chronological age is generally associated with the deviations in multiple health parameters that when measured can be combined into a clinical ‘frailty-score’^30^. As expected R-Atg5i mice displayed an initial increase in their frailty scores during autophagy inhibition in comparison to littermate controls, yet once mice have been switched back to a diet absent of dox, the frailty scores displayed a significant decrease over the next four months (Fig. 4c, Supplementary Movie. 1). In contrast, LT-Atg5i mice treated on dox for 6 months (median survival is around ~6 months on dox) continued to display a significant difference in their frailty scores, while almost all LT-Atg5i mice had already succumbed by eight-months (Fig. 4c). A similar increase in frailty was also noted in the LT-Atg5i_2 cohorts (Supplementary Fig. 4b). The penetrant kyphosis phenotype was largely irreversible, however 3/26 R-Atg5i mice did show evidence of recovery from kyphosis, while no mice displayed a reversal of the greying phenotypes. As such, while autophagy inhibition *in vivo* appears to promote frailty, autophagy restoration is seemingly able to substantially reverse this effect.

Remarkably the profound immune-associated phenotypes that we observed in autophagy-deficient LT-Atg5i mice were reversed in R-Atg5i mice. Serum markers of inflammation and WBC counts were indistinguishable between R-Atg5i and R-Control mice (Fig. 4d, e and Supplementary Fig. 5a). However, it should be noted that there was a trend towards a larger red blood cell distribution width (RDW) in aged R-Atg5i mice removed from dox for 8 months (14 months old), which has previously been linked to a range of diseases and an increased risk of acute myeloid leukemia (AML) (Fig. 4f)^31^. Additionally, R-Atg5i livers displayed a complete reversal of hepatomegaly and serum ALT levels (Supplementary Fig. 5c and c). The kidneys of R-Atg5i mice appeared to recover from autophagy inhibition and lacked evidence of sclerotic and enlarged glomeruli (Supplementary Fig. 5 d-f). Consistently, serum albumin levels displayed evidence of normalisation, although there was still a trend for reduced levels in R-Atg5i mice at the time point tested, suggesting that liver and/or kidney functions are largely recovered, if not completely (Supplementary Fig. 5g).

Similarly, the protein aggregation marker p62/SQSTM1 in the liver appeared much reduced in R-Atg5i mice in comparison to the LT-Atg5i mice, yet a small but substantial number of cells still exhibited a marked accumulation of p62 aggregation in R-Atg5i mice that had been off dox for four months (Fig. 5a). Additionally, R-Atg5i livers were also found to contain the presence of ceroid-laden macrophages and lipofuscin positivity, pigments known to increase with age and not seen in age-matched controls mice (Fig. 5b). Importantly, and in accordance with this partial restoration phenotype, molecular markers of aging such as TAF also remained significantly elevated in R-Atg5i mice. This is consistent with the persistent nature of telomeric DNA damage, which is reported to be irreparable^27, 32^. Together with other senescence markers (Fig. 4b), these data suggest that a portion of the cellular damage caused by a chronic block in autophagy is irreversible (Fig. 5c).

**Figure 5:**
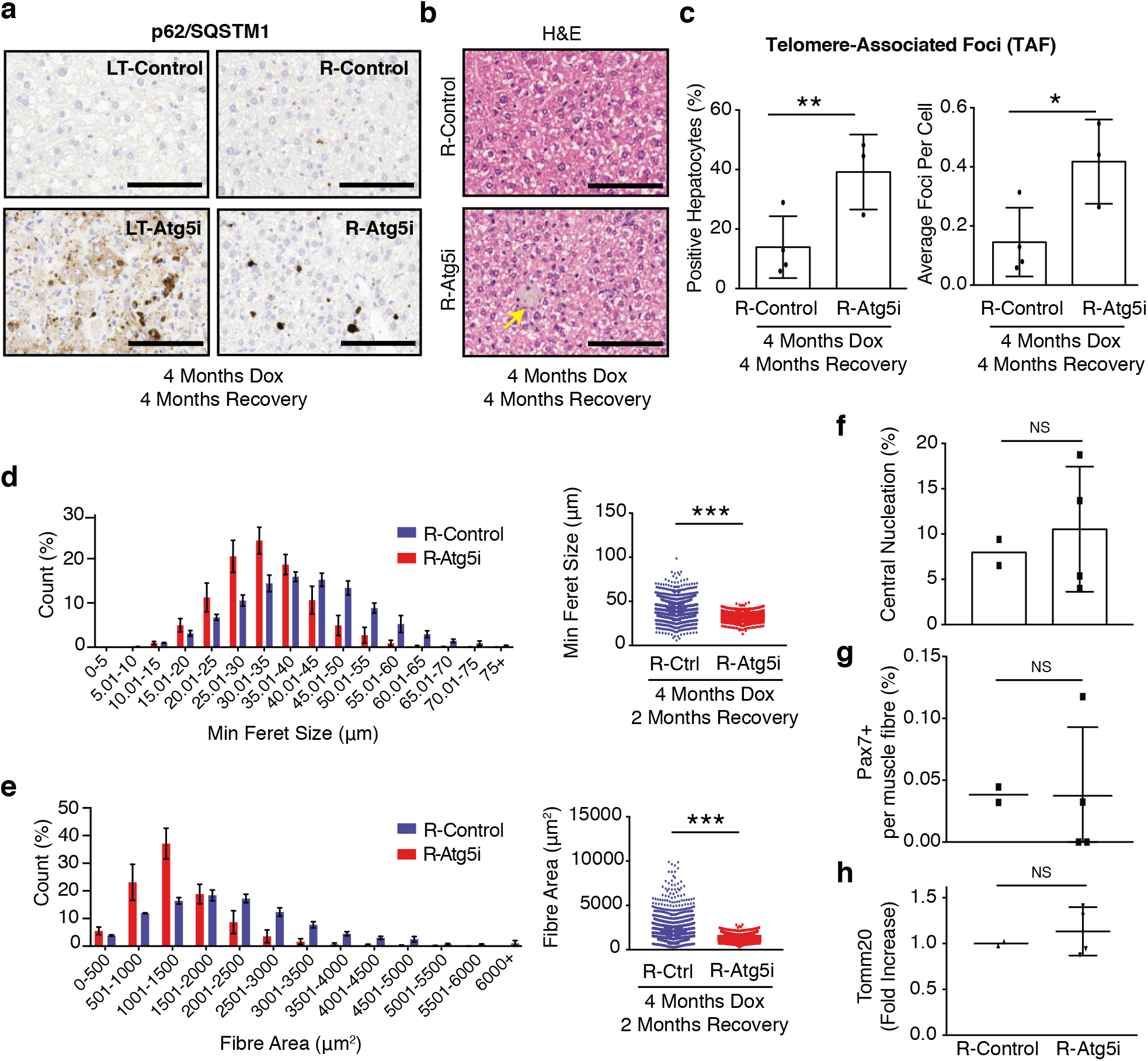
Restoration of autophagy does not reverse markers of aging. **a**, p62/Sqstm1 staining of R-Atg5i liver highlights the incomplete removal of aggregates four months after autophagy restoration. Scale bars,100 μm. **b**, The same livers have a higher incidence of age associated pigmentation in comparison to age-matched control mice. (yellow arrow). **c**, TAF frequency and abundance also remains elevated in R-Atg5i mice (unpaired two-tailed t-test; n= 4 R-Ctrl and 3 R-Atg5i). **d-h**, Skeletal muscle analysis from four months dox treated and two months restored R-Atg5i mice. R-Atg5i muscle fibres continue to display significant alterations in **(d)** minimum feret size (n= 4 R-Ctrl and 3 R-Atg5i, Mann Whitney test) and **(e)** crosssectional area (n= 4 R-Ctrl and 3 R-Atg5i, Mann Whitney test), but with a recovery of **(f)** central nucleation. **(g)** Pax7 nuclear positivity per fibre and **(h)** positivity for the mitochondrial marker TOM20 displays a heterogeneous recovery pattern in these mice, as measured by tissue immunofluorescence. (**f-h**, unpaired two-tailed Welches t-test; n= 2 R-Ctrl and 4 R-Atg5i). Error bars indicate standard deviations. *p<0.05; **p<0.01, ***p<0.001

Morphological analysis of skeletal muscle from R-Atg5i mice with autophagy restored suggests that muscle fibre size and morphology display no sign of recovery at the timepoint analysed (Fig. 5d, e and Supplementary Fig. 6a, b). However central nucleation and satellite cell frequency appeared to display a heterogeneous pattern, with evidence of recovery apparent in some individuals (Fig. 5f, g). As expected with Atg5 restoration, mitochondrial levels as determined by Tom20 positivity were restored to control levels (Fig. 5h). Additionally, the cardiac fibrosis observed LT-Atg5i mice appears to still be present four months post dox removal in R-Atg5i cohorts (Supplementary Fig. 6c). Together these data suggest that autophagy restoration may have tissue and pathology specific limitations in the capacity to recover from the tissue and cellular damage induced upon its inhibition. Crucially, whilst some tissues, such as the liver, appear to recover, they are still associated with age-associated pathologies at the molecular level.

## Accelerated tumor development in R-Atg5i mice

As R-Atg5i mice displayed some evidence of organismal rejuvenation and an increase in overall health, we sought to determine if autophagy restoration is able to reinstate natural longevity to the level seen in littermate control mice, or whether the damage accumulation impacting on lifespan was irreversible. Remarkably, while the life-span of R-Atg5i mice was significantly extended in comparison to LT-Atg5i mice (median survival 493 days versus 185 days since treatment began, respectively), it was also significantly shorter than the R-Control cohorts (Fig. 6a). In marked contrast to LT-Atg5i mice, the cause of death was predominantly associated with the development of tumors with an increased frequency and at earlier timepoints (Fig. 6b-d). Of note a whole-body mosaic Atg5 knockout mouse model has been previously reported to only develop liver adenomas but without any malignant tumors^33^. Together, our data suggest that a temporary period of autophagy inhibition may be enough to induce irreversible cellular damage, which might facilitate tumor development cooperatively with the restoration of autophagy.

**Figure 6:**
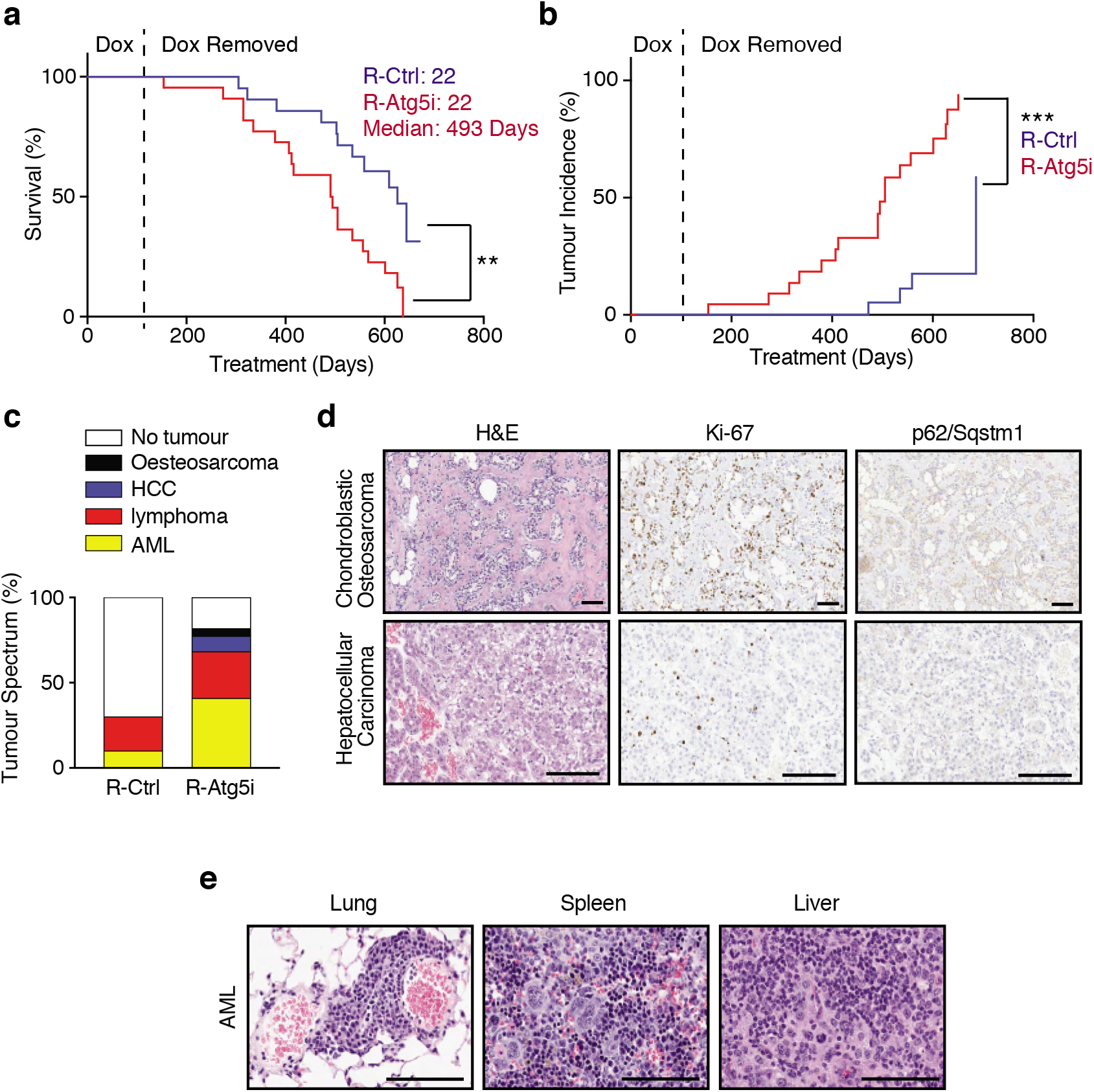
R-Atg5i mice are associated with accelerated spontaneous tumor development. **a**, R-Atg5i mice on display a reduced lifespan in comparison to R-Control mice (p<0.01). **b**, Increased frequency of spontaneous tumour formation in R-Atg5i cohorts (p<0.001). **c**, Tumor spectrum in R-Atg5i mice versus R-Control mice. **d – e**, Examples of R-Atg5i tumour histology. H&E staining and immunostaining of indicated proteins. Scale bars, 100μm.

## Discussion

While the rate of autophagic flux is believed to decrease with advancing age and has been postulated to be a driver of aging in multicellular organisms, evidence in mammals has been limited to the role of autophagy in maintaining stem cell populations^18, 21^. Such systemic organismal studies have been impossible to conduct owing to the embryonic or neonatal lethality, and rapid neurotoxicity in adult mice, that accompanies systemic autophagy ablation^14, 34^. The temporal control and lack of brain shRNA expression afforded by the Atg5i model have enabled us to circumvent these barriers, and separate developmental from tissue homeostatic effects that cannot be distinguished in aging models based on constitutive or *in utero* genetic modifications. Our findings support the theory that a reduction in autophagy is sufficient to induce several molecular and phenotypic characteristics associated with mammalian aging, including the development of age-associated diseases and a reduction in longevity. Here it is notable that our Atg5i mice phenocopy other models of aging driven by the accumulation of damage and in particular mitochondrial dysfunction^35, 36^.

Several health and life-span extending regimens in mammals, such as calorie restriction or pharmacological modulation, have been posited to exert their effects through the regulation of autophagy^7, 37^. However, these effects are also pleiotropic in nature and alter a multitude of cellular processes, making it impossible to deconvolute and ascribe the role of autophagy in these settings. Whilst recent genetic models that promote autophagic flux continuously throughout life have demonstrated an extension of health- and life-span in mammalian systems^11, 12^, it is unclear if the damage established by a loss of autophagy is sufficient for age acceleration and can be reversed. If therapeutic regimens in humans are to be established later in life, once autophagy-associated damage has accumulated, ascertaining the capacity for autophagy restoration to repair this damage is critical. In our model, systemic inflammation and frailty scores displayed a marked improvement upon autophagy restoration, which resulted in increased survival. However, while some tissues (i.e. liver and heart) displayed macroscopic normalisation, further analysis highlighted the persistence of pathological phenotypes. Our results indicate that the reversibility of markers of aging such as TAF, or macroscopic phenotypes such as greying and kyphosis may not recover. It should also be noted that we have chosen a late time-point to restore autophagy as this provided a clear and ubiquitous distinction between control and autophagy inhibited mice, shorter time points or intermittent dosing regimens may display further heterogeneity in damage and recovery phenotypes.

Our unexpected finding, that the temporal inhibition of autophagy predisposes to increased tumor development, provides a potential genetic explanation for the context-dependent role of autophagy in tumorigenesis^38, 39^: i.e. autophagy can be a tumor suppressor^33,40,41^ or a tumor promoter^42–44^. The irreversible damage induced by autophagy inhibition (e.g. genomic instability), might confer tumor susceptibility, while autophagy activity is perhaps required for actual malignant transformation. The clinical implication of our data is not limited to the advanced age state. As some pathophysiological states, such as obesity, are associated with an insufficient level of autophagy^45^, it would be interesting to determine if obese individuals retain an increased risk of tumor development even upon weight loss, in comparison to never obese populations.

## Supporting information

Supplemental Move 1

## Acknowledgements

We thank members of the Narita group, as well as K. Inoki of the University of Michigan, for their insights and suggestions. We are grateful to the following CRUK Cambridge Institute core facilities for advice and assistance: Histopathology, Light Microscopy (in particular H. Zecchini), and BRU.

## Author Contributions

L.D.C and M.N designed the research plan and interpreted the results. A.R.Y and C.N.J.Y isolated skeletal muscle tissue. C.N.J.Y performed staining and analysis of muscle sections. E.J.S and R.B are trained pathologists and reviewed all tissue slides. E.F and M.W established and assisted with the frailty scoring. K.A.W and M.C.H.C performed serum cytokine analyses. D.J and J.F.P performed the TAF studies. L.D.C and M.N wrote the manuscript, all authors viewed and commented on.

## Data and materials availability

All data and materials are available in the manuscript or upon request.

## Methods

### Atg5i mouse maintenance and aging

The generation and initial characterization of the Atg5i transgenic line has previously been described in detail^13^. Mice were maintained on a mixed C57Bl/6 × 129 background with littermate controls used in all experiments. All experimental mice were maintained as heterozygous for both the shRNA allele and CAG-rtTA3 alleles, whereas control littermates were lacking one of the alleles. Guide sequences were as follows: Atg5i TATGAAGAAAGTTATCTGGGTA^13^; Atg5i_2 TTATTTAAAAATCTCTCACTGT. Mice were maintained in a specific pathogen-free environment under a 12-h light/dark cycle, having free access to food and water. These mice were fed either a laboratory diet (PicoLab Mouse Diet 20, 5R58) or the same diet containing doxycycline at 200 ppm (PicoLab Mouse Diet, 5A5X). For this study mice were aged for two months before doxycycline administration in the diet. Mice were enrolled either to time-point study groups or long-term longevity cohorts (LT-and R-groups). Experienced animal technicians checked mice daily in a blinded fashion, and additionally mice were weighed and hand-checked on a weekly basis. Mice found to be of deteriorating health were culled under the advice of senior animal technicians if displaying end of life criteria. These signs include a combination of (1) hunched body position with matted fur, (2) piloerection, (3) poor body condition (BC) score (BC1 to 2), (4) failure to eat or drink, (5) cold to touch, and or (6) reduced mobility, including severe balance disturbances and ataxia. In accordance with UK home office regulations any mice suffering a 15% loss of body weight were also considered to be at an end-point. Note that for LT-longevity cohorts a portion of control mice were culled to generate age-matched littermate control tissue. These mice are marked as censored events on the survival curve. For analysis mice were treated as alive up to the point of their removal from the study where they are considered lost to follow-up and are not included in the calculations of median longevity. All experiments were performed in accordance with national and institutional guidelines, and the ethics review committee of the University of Cambridge approved this study.

### Frailty Scoring

Clinical frailty scoring was determined using the previously published frailty index^30^. A blinded researcher and animal technician performed all frailty scores independently within the same 48 hr period and scores were compared afterwards to ensure accuracy of phenotype scoring.

### Pathology and Immunohistochemistry

Explanted tissues were fixed in 10% neutral-buffered formalin solution for 24 hr and transferred to 70% ethanol. Tissues were embedded in paraffin, cut in 3μm sections on poly-lysine coated slides, deparaffinized, rehydrated, and stained with H&E. The PAS, Congo Red and Massons Trichrome histochemical stains were performed according to established protocols. An experienced pathologist reviewed all histology blinded for evidence of tumors and tissue pathologies. For immunohistochemistry and tissue immunoflourescence formalin-fixed paraffin-embedded samples were de-waxed and rehydrated. For anti-P21 (Santa Cruz, SC-6246; 1:500), and anti-TOM20 (Santa Cruz SC-11415, 1:500) staining antigen unmasking was performed with citrate buffer (10 mM sodium citrate, 0.05% Tween 20, pH 6) in a pressure cooker for 5 min at 120°C. For P21 exogenous peroxidases were quenched in 3% H2O2/PBS for 15 min and the remaining steps were performed according to Vector Labs Mouse on Mouse staining kit (MP-2400). The remaining antibodies were used at the following concentrations and ran on the Leica Polymer Detection system (DS9800) with the Leica automated Bond platform: Anti-SQSTM1 (Enzo, BML-PW9860; 1:750), anti-KI67 (Bathyl Laboratories, IHC-00375; 1:1000), Anti-LC3 (Nanotools, LC3-5F10 0231-100, 1:400).

For TOM20 analysis the intensity of signal per entire muscle section was determined and an average measurement of intensity per unit area calculated. Samples were then plotted as a fold increase relative to the average intensity per unit of control muscle sections For kidney glomeruli size tissue sections were analysed using ImageScope™ (Leica Biosystems) and the cross-sectional area of ten glomeruli in the renal cortex was reported per sample.

### Western Blotting

Western blot analysis was performed as previously^23^. Tissue samples were homogenized with the Precellys 24 tissue homogenizer in laemmeli buffer and samples ran on 12.5% or 15% gels. Protein was transferred to PVDF membranes (Immobilon, Millipore), which was subsequently blocked for 1 hr at room temperature (5% milk solution in TBS-Tween 0.1%) before incubating with primary antibody at 4°C overnight. An appropriate HRP-conjugated secondary antibody was incubated at room temperature for 1 hr. Western blots were visualized with chemiluminsence reagents (Sigma, RPN2106). Antibodies were used at the following concentrations: Anti-ATG5 (Abcam, ab108327; 1:1000), anti-LC3 (Abcam, ab192890; 1:1000), anti-ACTIN (Santa Cruz Biotechnology, I-19; 1:5000 [no longer commercially available]), anti-P53 (Cell Signalling Technologies, Clone 1C12; 1:1000), anti-P21 (Santa Cruz, SC-6246; 1:1000), anti-Histone H3 (Abcam, ab1791; 1:5000), anti-P16 (Santa Cruz, SC-1207; 1:1000), anti-HMGA1 (Abcam, ab129153; 1:1000).

### Blood and serum analysis

Whole blood composition was performed using the Mythic Hematology Analyser to determine whole blood counts, immune composition, and RDW. Mouse cytokines were determined using a cytometric bead array (BD Biosciences, Catalogue number: 552364). Sera isolated from mice were analyzed by the Core Biochemical Assay Laboratory (CBAL), Cambridge, UK for Alanine Transferase (Siemens Healthcare), Albumin (Siemens Healthcare), Bilirubin (Siemens Healthcare), and Creatinine (Siemens Healthcare) using automated Siemens Dimension RxL and ExL analyzers.

### Telomere Associated DNA Damage Foci (TAF)

Formalin-fixed paraffin-embedded liver sections were hydrated by incubation in 100% Histoclear, 100, 95 and 2X 70% methanol for 5 min before washed in distilled water for 2X 5 min. For antigen retrieval, the slides were placed in 0.1 M citrate buffer and heated until boiling for 10 min. After cooling down to room temperature, the slides were washed 2X with distilled water for 5 min. After blocking in normal goat serum (1:60) in BSA/PBS, anti-γ-H2A.X primary antibody (Cell Signalling Technologies, S139; 1:250) was applied and incubated at 4 °C overnight. Slides were washed 3X in PBS, incubated with secondary antibody for 30 min, washed three times in PBS and incubated with Avidin DCS (1:500) for 20 min. Following incubation, slides were washed three times in PBS and dehydrated with 70, 90 and 100% ethanol for 3 min each. Sections were denatured for 5 min at 80 °C in hybridization buffer (70% formamide (Sigma), 25 mM MgCl_2_, 1 M Tris pH 7.2, 5% blocking reagent (Roche) containing 2.5 μg ml^-1^ Cy-3-labelled telomere specific (CCCTAA) peptide nuclei acid probe (Panagene), followed by hybridization for 2 h at room temperature in the dark. The slides were washed with 70% formamide in 2×SSC for 2X 15 min, followed by 2×SSC and PBS washes for 10 min. Sections were incubated with DAPI, mounted and imaged. In depth Z stacking was used (a minimum of 40 optical slices with ×100 objective) followed by Huygens (SVI) deconvolution.

### Senescence associated beta-galactosidase staining

Whole tissue samples were washed in PBS (pH5.5) before being fixed in 0.5% Glutaraldehyde overnight and washed 2X 15 min in PBS (pH5.5) at 4^°^C. SA-β-gal activity was assessed after incubation in X-Gal solution for 90 minutes at 37^°^C.

### Muscle Morphopmetric Analysis

Mice were sacrificed at the time points described and dissected muscle was rapidly frozen in liquid nitrogen cooled isopentane to maintain structure and minimize tissue artifacts. Experimental mice and age-matched littermate controls were isolated at the same time to ensure processing was consistent between groups. Frozen muscles were equilibrated in a cryostat chamber to −20°C and cryosections 10-μm thick were then cut from the middle third of the sample and collected on poly-L-lysine (0.5 mg/ml)–coated glass slides. Sections were allowed to air dry and were then frozen at −80°C prior to use. Samples were brought to 4°C on ice and fixed in a 4% w/v 0.45 mm filtered paraformaldehyde solution in 1 × PBS for 15 min at 4°C. PFA was removed by three 5 min washes in 1 × PBS, then blocked in 10% v/v serum in 1 × PBST (0.01% Tween-20) for 1 hr at RT. Primary anti-dystrophin antibody (Abcam, ab15277, 1:1000) was then applied in 1 × PBST containing 10% v/v serum for 2 h at room temperature. Three 5-min PBST washes were applied before secondary antibody conjugated to Alexa Fluor 647, with DAPI at 1:1000, incubation in PBST and 10% v/v serum for 1 h at room temperature. Sections were finally washed three times for 15 min before mounting in Vectorshield Antifade Mounting Medium (Vector Labs). Whole cross-sections of TA muscles were produced via montaged 40× magnification tile scans (Zeiss Axio Z1 Widefield system). Morphometric analysis was performed using Fiji open source software as previously described (26461208). Simultaneous DAPI nuclear stain was used for central nucleation count. PAX7 counts were performed manually in a blinded fashion, a satellite cell was defined as having a PAX7 positive nuclei within a LAMININ cell border staining. For immunostaining the following antibodies were used anti-PAX7 PAX7 (DSHB, PAX7, 1:50), after pretreatment with Vector Labs Mouse on Mouse Blocking Reagent (MKB-2213) according to manufacturer’s instructions and anti-LAMININ (Abcam, ab11576, 1:1000).

---

**Supplementary Figure 1:**
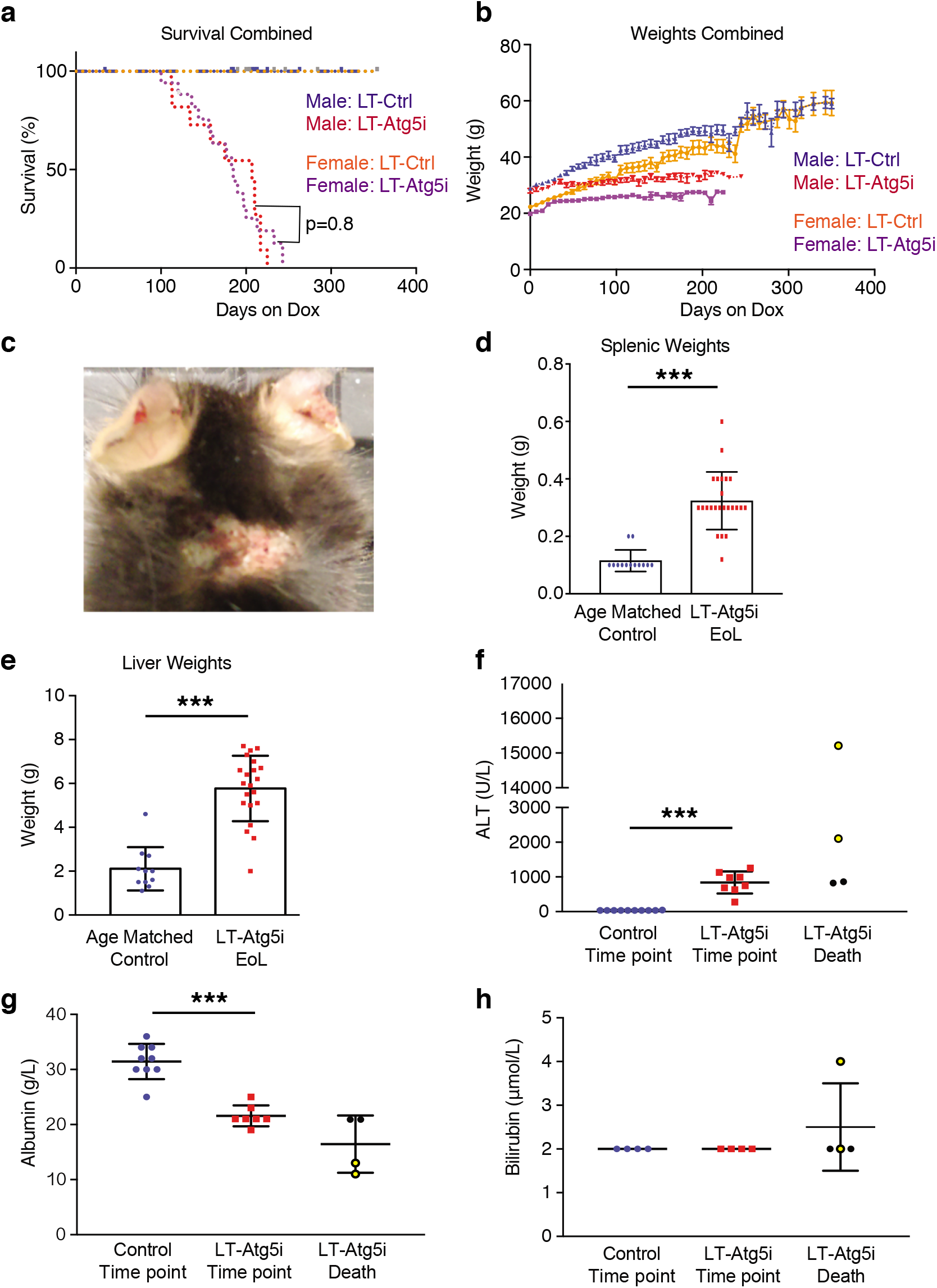
Characterisation of LT-Atg5i mice. **a**, LT-Atg5i mice display no life-span associated sex bias (Red, LT-Atg5i Males; Purple, LT-Atg5i Females; p=0.8). **b**, LT-Atg5i mouse weight plateau while LT-Control mice continue to gain weight over their lifetime. **c**, Example of mouse suffering from ulcerative dermatitis. **d**, Splenic weights were increased in LT-Atg5i mice in comparison to age matched LT-Control mice. **e**, LT-Atg5i mice also display an increase in liver weight. **f-h**, liver function of LT-Atg5i mice as determined using serum samples. LT-Atg5i mice on dox for 4 months display (**f**) an increase in serum ALT and (**g**) a decrease in serum albumin that is further exacerbated in a subset of LT-Atg5i EoL individuals (yellow circles). (**h**) The only sample tested that displayed an increase in serum bilirubin levels was also from a mouse displaying high levels of serum ALT and low levels of serum album. Error bars indicate standard deviations. *p<0.05; **p<0.01, ***p<0.001

**Supplementary Figure 2:**
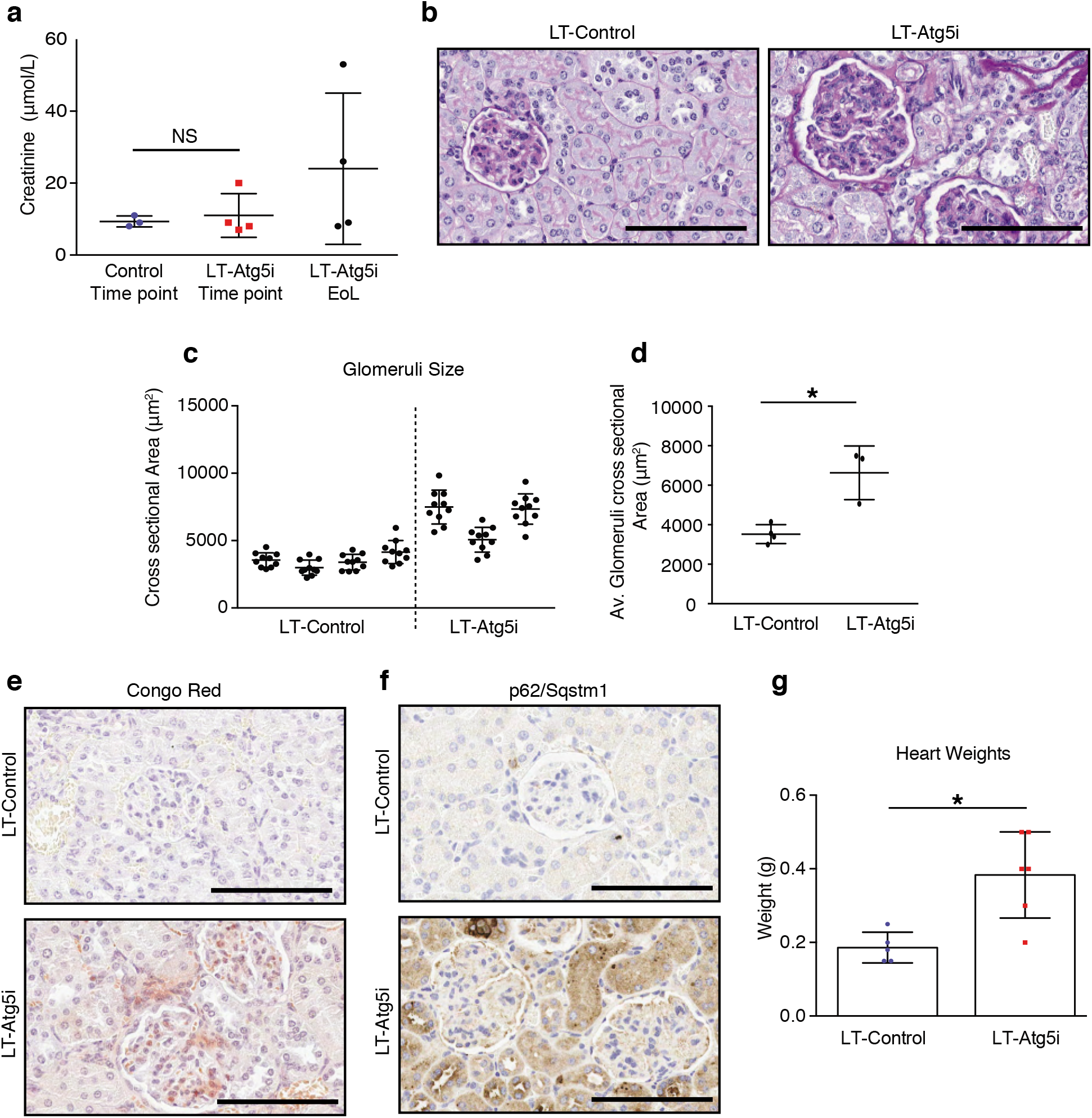
Kidney alterations in LT-Atg5i mice. (**a**) LT-cohorts treated with doxycycline for 6 months mice display no significant differences in serum creatinine levels (unpaired two-tailed Welches t-test, NS denotes not significant; n= 3 LT-Control and 4 LT-Atg5i). At death only a subset of LT-Atg5i mice display an increase in serum creatinine levels. **b-f**, LT-Atg5i mouse kidneys treated with doxycycline for 6 months present with (**b**) evidence of sclerotic glomeruli determined using PAS stain that are also (**c-d**) enlarged and hypercelluar in comparison to LT-Control (p=0.0479, unpaired two-tailed t-test; n= 4 LT-Control and 3 LT-Atg5i, the cross-sectional area of 10 randomly chosen glomeruli were measured per mouse). (**e**) Congo red and (**f**) P62/Sqstm1 staining of LT-Atg5i mouse kidneys treated with doxycycline for 6 months highlights an increase in protein aggregation not present in age-matched LT-Control mice. **g**, Cardiac tissue from LT-Atg5i mice at death was significantly heavier than age-matched LT-Control mice. (p=0.0108). Error bars indicate standard deviations. *p<0.05; **p<0.01, ***p<0.001

**Supplementary Figure 3:**
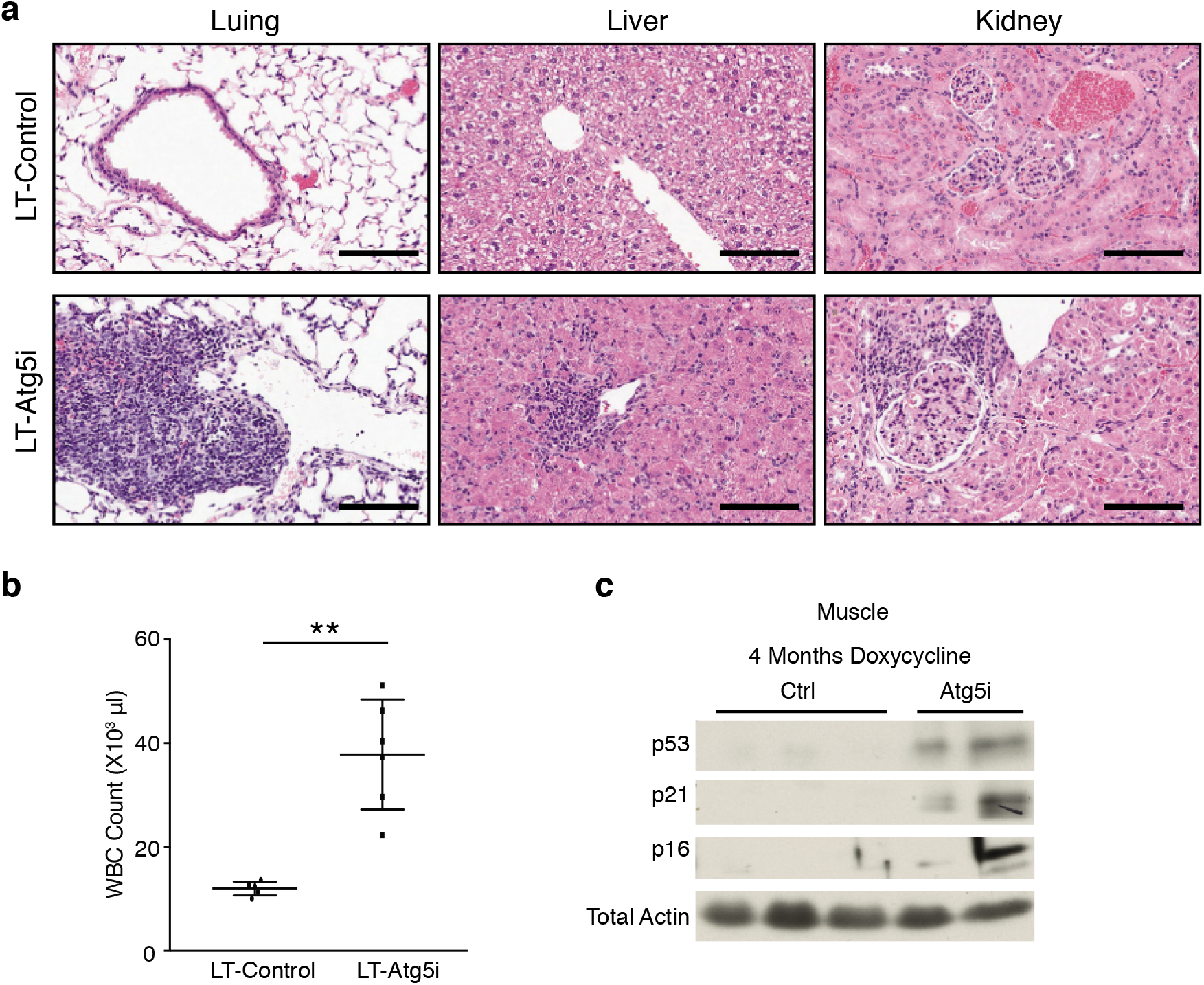
Systemic alterations in LT-Atg5i mice. **a**, LT-Atg5i mice display evidence of widespread immune infiltration across multiple tissues in comparison to age-matched controls. Scale bars,100 μm. **b**, White blood cell counts (WBC) of LT-Control and LT-Atg5i mice treated with doxycycline for 4 months (6 months old) (unpaired two-tailed Welches t-test, n=5-6 per group). **c**, Skeletal muscle displays markers of senescence in LT-Atg5i cohorts on doxycycline for 4 months. **p<0.01

**Supplementary Figure 4:**
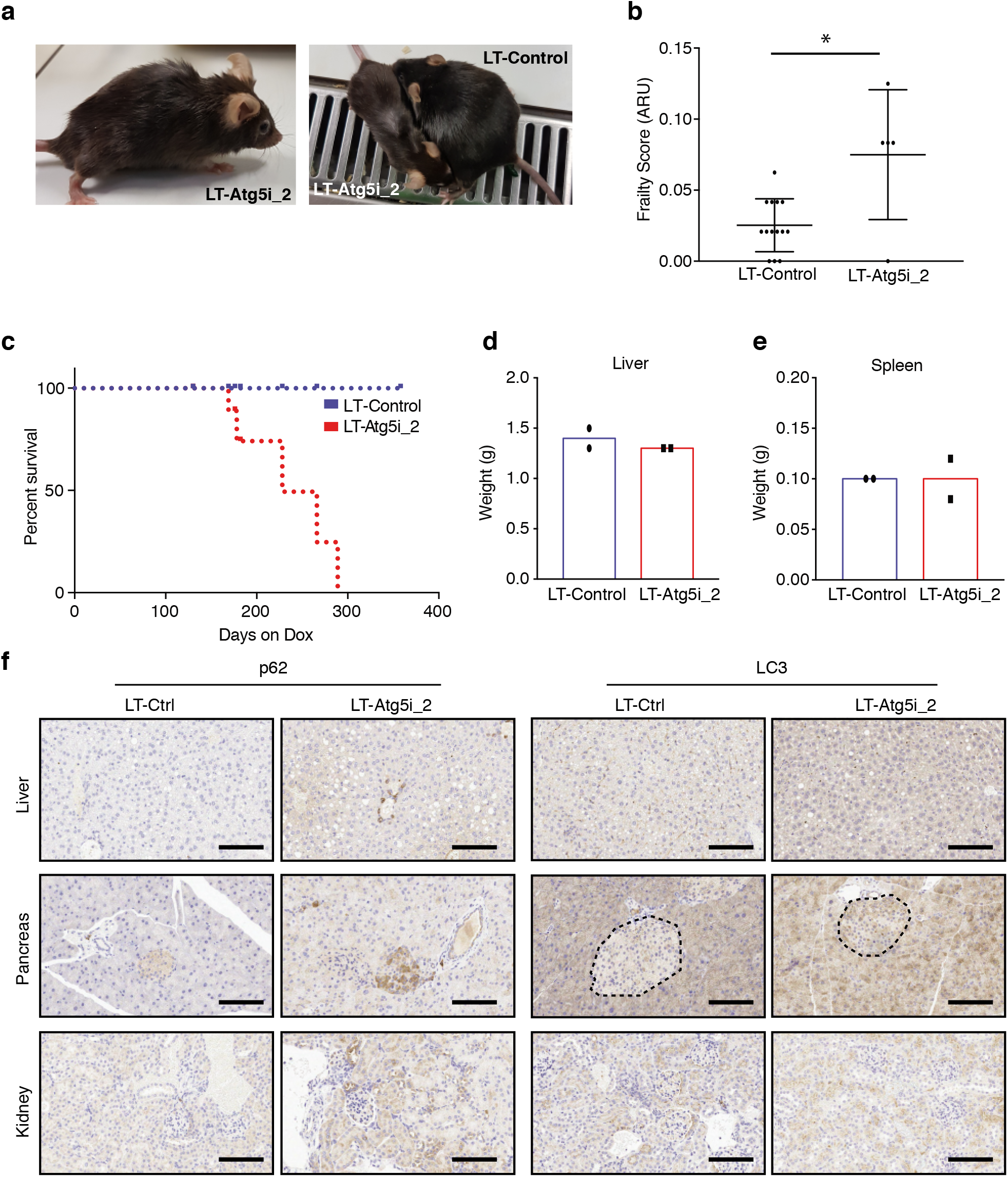
Hypomorphic LT-Atg5i_2 mice also display aging phenotypes. **a-c**, LT-Atg5i_2 mice phenotypically recapitulate premature ageing phenotypes including (**a**) kyphosis, (**b**) increased frailty (ARU, arbitrary units; Mann-whitney n= 14 LT-Control and 5 LT-Atg5i_2 mice), and (**c**) reduced longevity. **d-f** However, Atg5i_2 mice appear to have a hypomorphic phenotype and do not recapitulate the phenotypes found in Atg5 knock-out and LT-Atg5i. These include no evidence of (d) hepatomegaly or (**e**) splenomegaly. (**f**) Correspondingly, p62/SQSTM1 and LC3 levels do not accumulate to the same degree in LT-Atg5_2 mice treated with doxycycline for 6 weeks. Scale bars,100 μm. Error bars indicate standard deviations. *p<0.05

**Supplementary Figure 5:**
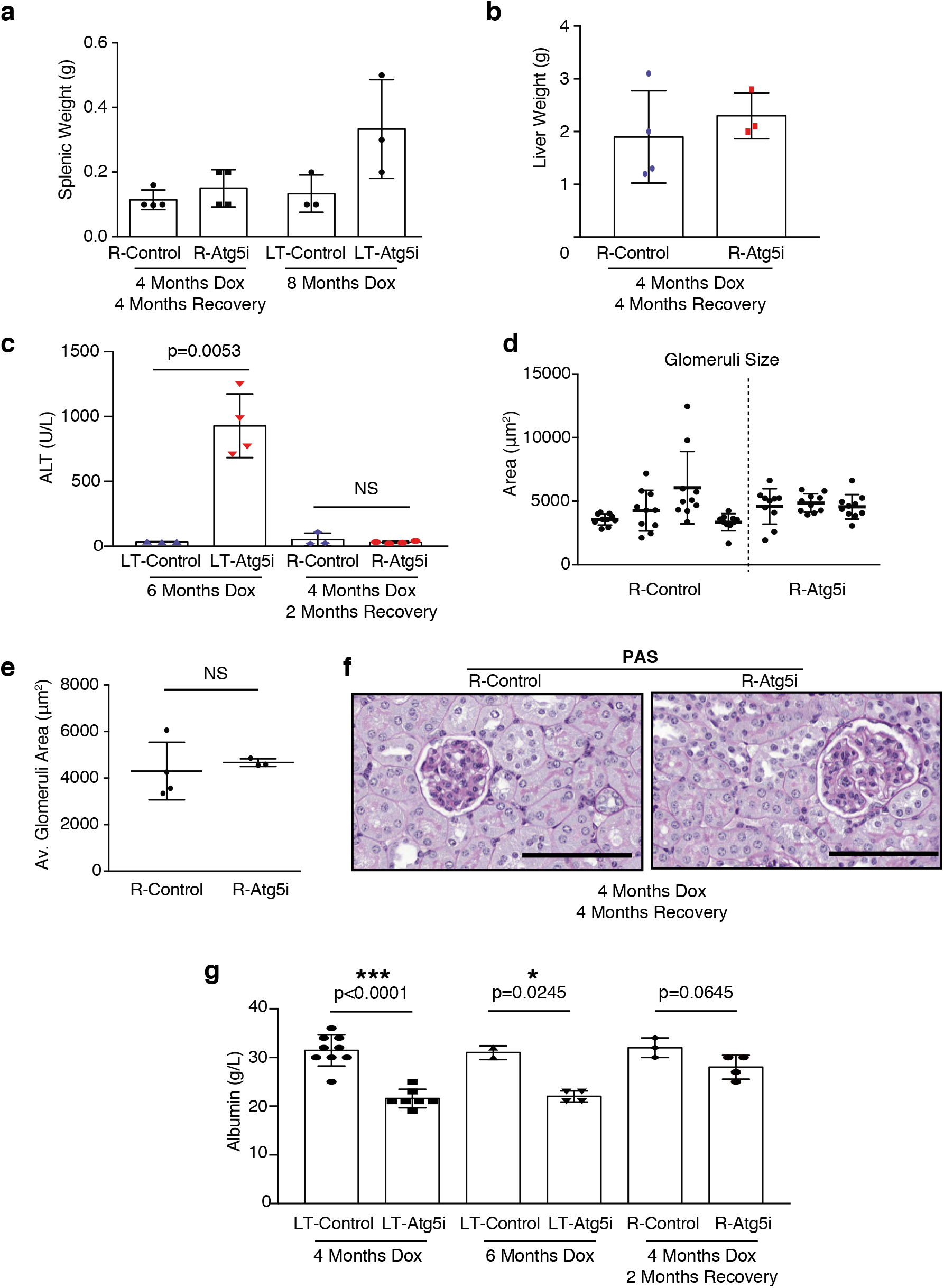
Autophagy restoration reverses hepatomegaly and splenomegaly. **a**, Splenic and **b**, liver weights from R-Atg5i mice exhibit evidence of recovery. **c**, In addition R-Atg5i mice display a reduction in serum ALT levels (unpaired two-tailed Welches t-test; n= 3-4 per cohort). **d-f**, R-Atg5i mice 4 months post dox removal display evidence of recovery in the kidneys as determined by (**d-e**) normalisation of glomeruli size appeared relative to age-matched controls (unpaired two-tailed Mann whitney, n=3-4 mice per group) and (**f**) the absence of sclerosis. **g**, A partial recovery in serum albumin levels is also present in these mice unpaired two-tailed Welches t-test; n= 2-9 per cohort). Error bars indicate standard deviations. *p<0.05; **p<0.01, ***p<0.001

**Supplementary Figure 6:**
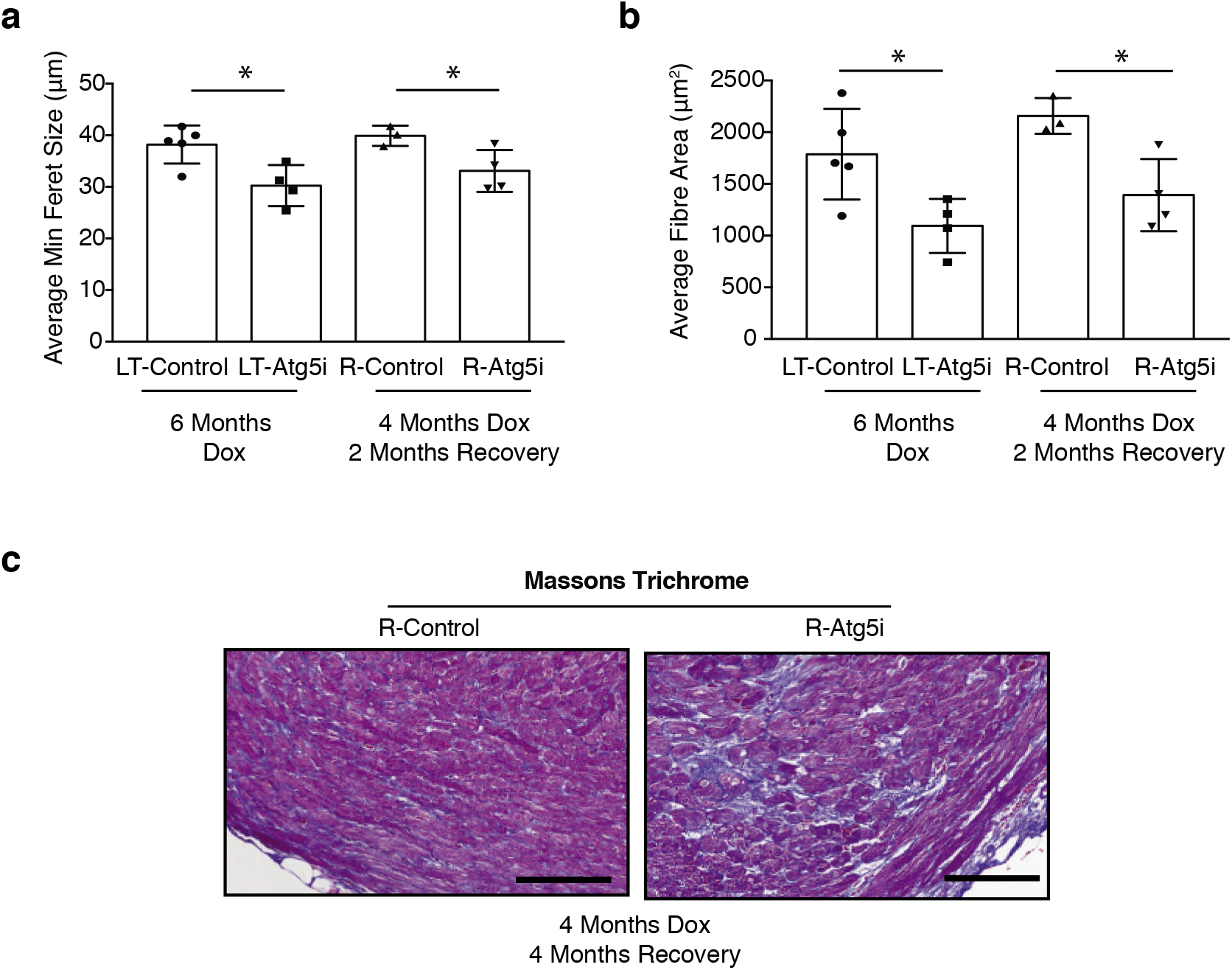
Autophagy restoration displays segmental rescue of tissue phenotypes. **a-b**, Skeletal muscle displays no rescue of phenotype once Atg5i mice are removed from dox. As determined by (**a**) minimal feret size, and (**b**) cross-sectional area. (unpaired two-tailed Welches t-test, n=3-5 per group). **c**, Cardiac fibrosis was still present in R-Atg5i mice 4 months post dox removal. Error bars indicate standard deviations. *p<0.05

**Supplementary Movie. 1**: R-Atg5i mice 4 months post dox removal highlighting the stochastic response to autophagy restoration. All mice were treated on dox for 4 months before dox removal for 2 months. At this stage, 100 % of mice show kyphosis. The movie represents three examples with different levels of recovery. Mouse I exhibits little recovery, whereas mouse III looks normal with no sign of kyphosis. Mouse II appears active but with mild kyphosis.

## References and Notes

1. Partridge, L., Deelen, J. & Slagboom, P. E. Facing up to the global challenges of ageing. Nature 561, 45–56 (2018).

2. Sen, P., Shah, P. P., Nativio, R. & Berger, S. L. Epigenetic Mechanisms of Longevity and Aging. Cell 166, 822–839 (2016).

3. López-Otín, C., Galluzzi, L., Freije, J. M. P., Madeo, F. & Kroemer, G. Metabolic Control of Longevity. Cell 166, 802–821 (2016).

4. Newman, J. C. et al. Strategies and Challenges in Clinical Trials Targeting Human Aging. J. Gerontol. A Biol. Sci. Med. Sci. 71, 1424–1434 (2016).

5. Colman, R. J. et al. Caloric restriction reduces age-related and all-cause mortality in rhesus monkeys. Nat Commun 5, 3557 (2014).

6. Mattison, J. A. et al. Impact of caloric restriction on health and survival in rhesus monkeys from the NIA study. Nature 489, 318–321 (2012).

7. Rubinsztein, D. C., Mariño, G. & Kroemer, G. Autophagy and Aging. Cell 146, 682–695 (2011).

8. Meléndez, A. et al. Autophagy genes are essential for dauer development and life-span extension in C. elegans. Science 301, 1387–1391 (2003).

9. Bjedov, I. et al. Mechanisms of life span extension by rapamycin in the fruit fly Drosophila melanogaster. Cell Metab. 11, 35–46 (2010).

10. Jia, K. & Levine, B. Autophagy is required for dietary restriction-mediated life span extension in C. elegans. Autophagy 3, 597–599 (2007).

11. Fernández, Á. F. et al. Disruption of the beclin 1-BCL2 autophagy regulatory complex promotes longevity in mice. Nature 125, 85 (2018).

12. Pyo, J.-O. et al. Overexpression of Atg5 in mice activates autophagy and extends lifespan. Nat Commun 4, 2300 (2013).

13. Cassidy, L. D. et al. A novel Atg5-shRNA mouse model enables temporal control of Autophagy in vivo. Autophagy 1–11 (2018). doi:10.1080/15548627.2018.1458172

14. Komatsu, M. et al. Loss of autophagy in the central nervous system causes neurodegeneration in mice. Nature 441, 880–884 (2006).

15. Menzies, F. M., Fleming, A. & Rubinsztein, D. C. Compromised autophagy and neurodegenerative diseases. Nat. Rev. Neurosci. 16, 345–357 (2015).

16. Pettan-Brewer, C. & Treuting, P. M. Practical pathology of aging mice. Pathobiol Aging Age Relat Dis 1, 393 (2011).

17. Komatsu, M. et al. Impairment of starvation-induced and constitutive autophagy in Atg7-deficient mice. J Cell Biol 169, 425–434 (2005).

18. Ho, T. T. et al. Autophagy maintains the metabolism and function of young and old stem cells. Nature 543, 205–210 (2017).

19. Mortensen, M. et al. Loss of autophagy in erythroid cells leads to defective removal of mitochondria and severe anemia in vivo. Proc. Natl. Acad. Sci. U.S.A. 107, 832–837 (2010).

20. Watson, A. S. et al. Autophagy limits proliferation and glycolytic metabolism in acute myeloid leukemia. Cell Death Discov 1, 371 (2015).

21. García-Prat, L. et al. Autophagy maintains stemness by preventing senescence. Nature 529, 37–42 (2016).

22. Sousa-Victor, P. et al. Geriatric muscle stem cells switch reversible quiescence into senescence. Nature 506, 316–321 (2014).

23. Young, A. R. J. et al. Autophagy mediates the mitotic senescence transition. Genes Dev. 23, 798–803 (2009).

24. Narita, M. et al. Spatial coupling of mTOR and autophagy augments secretory phenotypes. Science 332, 966–970 (2011).

25. Kang, H. T., Lee, K. B., Kim, S. Y., Choi, H. R. & Park, S. C. Autophagy impairment induces premature senescence in primary human fibroblasts. PLoS ONE 6, e23367 (2011).

26. Dörr, J. R. et al. Synthetic lethal metabolic targeting of cellular senescence in cancer therapy. Nature 501, 421–425 (2013).

27. Hewitt, G. et al. Telomeres are favoured targets of a persistent DNA damage response in ageing and stress-induced senescence. Nat Commun 3, 708 (2012).

28. Correia-Melo, C. et al. Mitochondria are required for pro-ageing features of the senescent phenotype. EMBO J. 35, 724–742 (2016).

29. Jurk, D. et al. Chronic inflammation induces telomere dysfunction and accelerates ageing in mice. Nat Commun 2, 4172 (2014).

30. Whitehead, J. C. et al. A clinical frailty index in aging mice: comparisons with frailty index data in humans. J. Gerontol. A Biol. Sci. Med. Sci. 69, 621–632 (2014).

31. Abelson, S. et al. Prediction of acute myeloid leukaemia risk in healthy individuals. Nature 559, 400–404 (2018).

32. Fumagalli, M. et al. Telomeric DNA damage is irreparable and causes persistent DNA-damage-response activation. Nat. Cell Biol. 14, 355–365 (2012).

33. Takamura, A. et al. Autophagy-deficient mice develop multiple liver tumors. Genes Dev. 25, 795–800 (2011).

34. Karsli-Uzunbas, G. et al. Autophagy is required for glucose homeostasis and lung tumor maintenance. Cancer Discov 4, 914–927 (2014).

35. Trifunovic, A. et al. Premature ageing in mice expressing defective mitochondrial DNA polymerase. Nature 429, 417–423 (2004).

36. Kujoth, G. C. et al. Mitochondrial DNA mutations, oxidative stress, and apoptosis in mammalian aging. Science 309, 481–484 (2005).

37. Hansen, M., Rubinsztein, D. C. & Walker, D. W. Autophagy as a promoter of longevity: insights from model organisms. Nat. Rev. Mol. Cell Biol. 19, 579–593 (2018).

38. Guo, J. Y., Xia, B. & White, E. Autophagy-mediated tumor promotion. 155, 1216–1219 (2013).

39. Rosenfeldt, M. T. et al. p53 status determines the role of autophagy in pancreatic tumour development. Nature 504, 296–300 (2013).

40. Qu, X. et al. Promotion of tumorigenesis by heterozygous disruption of the beclin 1 autophagy gene. 112, 1809–1820 (2003).

41. Yue, Z., Jin, S., Yang, C., Levine, A. J. & Heintz, N. Beclin 1, an autophagy gene essential for early embryonic development, is a haploinsufficient tumor suppressor. Proc. Natl. Acad. Sci. U.S.A. 100, 15077–15082 (2003).

42. Guo, J. Y. et al. Autophagy suppresses progression of K-ras-induced lung tumors to oncocytomas and maintains lipid homeostasis. Genes Dev. 27, 1447–1461 (2013).

43. Strohecker, A. M. et al. Autophagy sustains mitochondrial glutamine metabolism and growth of BrafV600E-driven lung tumors. Cancer Discov 3, 12721–285 (2013).

44. Yang, A. et al. Autophagy Sustains Pancreatic Cancer Growth through Both Cell-Autonomous and Nonautonomous Mechanisms. Cancer Discov 8, 276–287 (2018).

45. Lim, Y. M. et al. Systemic autophagy insufficiency compromises adaptation to metabolic stress and facilitates progression from obesity to diabetes. Nat Commun 5, 4934 (2014).

